# DNA Polymerase Locks Replication Fork Under Stress

**DOI:** 10.1101/2024.10.09.617451

**Authors:** Xiaomeng Jia, James T. Inman, Anupam Singh, Smita S. Patel, Michelle D. Wang

## Abstract

Replication of DNA requires the parental DNA to be unwound to allow the genetic information to be faithfully duplicated by the replisome. While this function is usually shared by a host of proteins in the replisome, notably DNA polymerase (DNAP) and helicase, the consequence of DNAP synthesizing DNA while decoupled from helicase remains not well understood. The unwinding of downstream DNA poses significant stress to DNAP, and the interaction between DNAP and the replication fork may affect replication restart. In this work, we examined the consequences of DNAP working against the stress of the DNA replication fork. We found that prolonged exposure of DNAP to the stress of the replication fork inactivates replication. Surprisingly, replication inactivation was often accompanied by a strong DNAP interaction with the leading and lagging strands at the fork, locking the fork in place. We demonstrated that fork locking is a consequence of DNAP forward translocation, and the exonuclease activity of DNAP, which allows DNAP to move in reverse, is essential in protecting the fork from inactivation. Furthermore, we found the locking configuration is not reversible by the subsequent addition of helicase. Collectively, this study provides a deeper understanding of the DNAP-fork interaction and mechanism in keeping the replication fork active during replication stress.

---

DNA synthesis during replication is a highly dynamic process carried out by DNA polymerase (DNAP) in coordination with several other components within the replisome, such as helicase. Ideally, DNAP and helicase move synchronously at the replication fork, with helicase unwinding the DNA at the fork to allow DNAP access to the leading strand for replication. However, sometimes, this synchronization is disrupted. For example, if helicase lags behind DNAP or temporarily dissociates from the fork, the two separated DNA strands can reanneal, leading DNAP to work against the fork on its own. DNAP must overcome the physical barriers of unwinding the fork to continue synthesizing the new DNA strand. This leads to replication stress, which can pose significant challenges to the replication machinery.

Understanding how DNAP responds to these conditions is crucial for elucidating the mechanisms maintaining genomic stability during DNA replication.

The ability of DNAP to function independently of helicase under these conditions remains an important question that is not fully understood. The T7 replisome is a model system for mechanistic understanding of replication^1-4^ and serves as a simple model system to address this question. Previous studies show that when T7 DNAP encounters a DNA fork, it may replicate a few base pairs before stalling under the reannealing stress of the DNA in its front, which pushes the DNAP to move in reverse^5, 6^. DNAP can backtrack by removing the replicated DNA via its exonuclease activity, releasing the replication stress, which may provide a new opportunity for DNAP to forward translocate to replicate DNA. Thus, DNAP can shuttle between active replication and exonuclease activity during such stalling^7^. This stalling behavior raises questions about the enzyme’s efficiency in restarting replication once replication stress is alleviated. However, little is known about how stalling impacts the ability of DNAP to restart due to a lack of complete understanding of the nature of DNAP’s interaction with the replication fork.

This work aims to explore T7 DNAP’s interactions with the replication fork and investigate the impact of replication stress on the enzyme’s ability to resume DNA synthesis. We discovered a surprising behavior of DNAP’s interaction with the fork. When DNAP works at the fork on its own for a prolonged duration, it can lock the fork by interacting with both the leading and lagging strands. Once locked, the fork cannot be unlocked readily as DNAP binds tightly to the fork, even after helicase addition. Fork-locking results from DNAP synthesizing DNA while advancing the fork without helicase. Interestingly, the exonuclease activity of DNAP is essential to minimizing fork locking, revealing a new role of exonuclease in maintaining an active fork. These results on how the basal replication machinery DNAP interacts with a replication fork may have significant implications for understanding replication stalling and subsequent replication restart.

## Results

### DNAP can lock the replication fork

If T7 DNAP interacts with a replication fork for an extended duration, DNAP cannot replicate a substantial distance as the fork represents a significant resistance to DNA replication. This raises the possibility that this resistance may inactivate the bound DNAP. To investigate this possibility, we employed two techniques that we previously developed^8-11^ to study protein-DNA interactions via mechanical unzipping of a DNA molecule using an optical trap (Fig. 1a, Fig. S1b). The “unzipping tracker” uses the DNA fork to track the trajectory of a translocating motor in real-time, and the “unzipping mapper” detects the location and strength of a bound protein via rapid mechanical unzipping through the bound protein.

**Figure 1.**
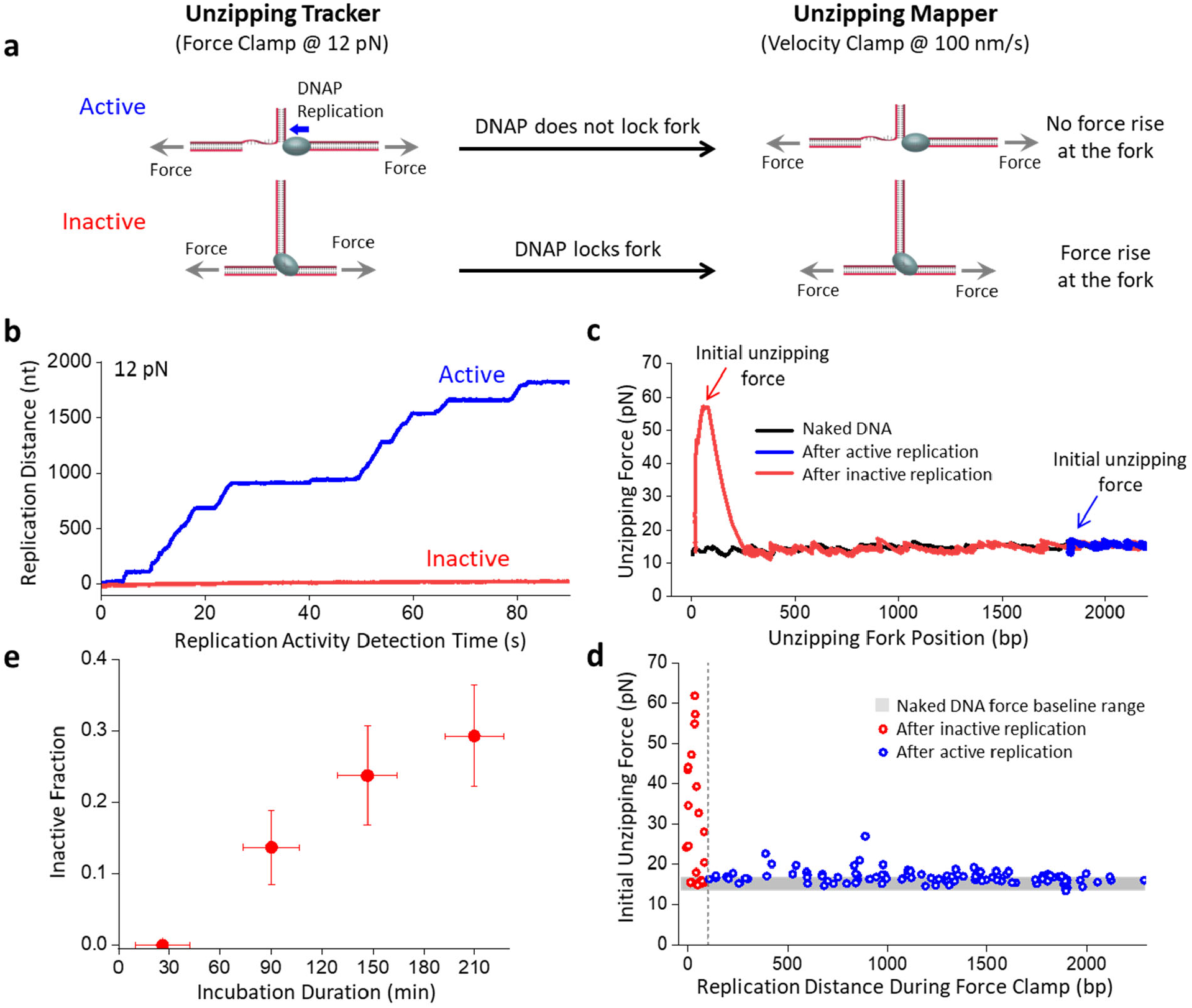
DNAP can lock the replication fork. **a**.Experimental techniques for investigating DNAP interactions with replication fork. First, the unzipping tracker uses the DNA fork to track the trajectory of the DNAP in real-time. Active forks held under a constant force result in an increase in the tether extension while inactive forks do not increase the extension. Subsequently, the unzipping mapper detects the location and strength of a bound protein interacting with the fork via mechanical unzipping through the remaining DNA. A locked fork results in a force rise at the start of the unzipping mapper step while an unlocked fork has no detectable force rise. **b**. Representative active (blue) and inactive (red) traces during the unzipping tracker step. The replication distance of the DNAP is tracked for 90 s before proceeding to the unzipping mapper step. **c**. Immediately after the unzipping tracker step, the unzipping mapper is employed on the same traces shown in **b** to detect any interactions of DNAP with the remaining parental DNA. The unzipping fork position at the start of the unzipping mapper corresponds to the fork position after replication in the unzipping tracker step. At this initial unzipping fork position, the initial unzipping force indicates resistance to strand opening for the inactive trace (red) and minimal resistance for the active trace (blue). The unzipping mapper curve of naked DNA in the absence of the DNAP is shown in black. **d**. The initial unzipping force from the beginning of the unzipping mapper versus the replication distance during the unzipping tracker from *N* = 137 individual traces. Traces identified as active during the replication tracker are shown in blue, and traces identified as inactive in red. **e**. The inactive fraction versus incubation duration of DNAP with DNA forks. Error bars represent the standard deviation, with *N* = 41, 44, 38, and 41 individual traces for 26, 90, 147, and 210 min incubation, respectively.

In this experiment, we first incubated replication forks (Fig. S1a) with T7 DNAP for a specified duration. Subsequently, we used the unzipping tracker to examine whether a replication fork was still active by holding the fork at a force of 12 pN, which facilitates DNA unzipping and allow leading strand DNA synthesis without helicase^3, 4, 12^ (Fig. 1a,b). If a trace showed active replication, DNA extension increased by 0.77 nm for each nucleotide replicated (Methods). In contrast, inactive replication did not increase DNA extension.

Immediately following this step, we employed the unzipping mapper and rapidly unzipped through the remainder of the parental DNA to detect any interactions of DNAP with the parental DNA (Fig. 1a,c). We noticed that an active trace typically showed no force rise above the naked DNA baseline during the unzipping mapper step. Surprisingly, an inactive trace often showed a force rise significantly above the naked DNA baseline at the beginning of the unzipping. This force rise reveals that the fork experiences resistance to strand opening. Since such resistance was not present without DNAP, this finding suggests that a DNAP can simultaneously interact with both the leading and lagging strands at the fork, “locking” the fork and rendering it inactive.

We found a strong correlation between a fork’s activity during the unzipping tracker step and the subsequent unzipping force rise during the unzipping mapper step: an inactive fork typically leads to a force rise (Fig. 1d). The average disruption force is about 27.2 pN, comparable to forces required to disrupt a bound *E. coli* RNA polymerase^13, 14^, a Cas protein^10^, restriction enzymes^11, 15, 16^, or a nucleosome^17-20^, which can also be significant barriers to replication^21-23^. Furthermore, we found that forks became increasingly less active with an increase in the incubation duration (Fig. 1e). To exclude the possibility that fork locking is a consequence of DNAP degradation at room temperature over the incubation period, we left the DNAP in a test tube without any DNA fork for 4 hours and tested its activity. We found that the DNAP activity remained unchanged (Fig.S2a). Thus, these data demonstrate that prolonged interaction of DNAP with a replication fork leads to DNAP locking and inactivation of the fork.

### DNAP exonuclease activity limits fork-locking

Since T7 DNAP can support both polymerization activity and a 3’ to 5’ exonuclease activity, we first examined how the exonuclease activity impacted the observed DNAP fork-locking behavior. Thus, we repeated the experiments shown in Fig. 1 using two DNAP mutants lacking exonuclease activity (Fig. 2, Fig.S3): one containing two point-mutations, which we refer to as exo-DNAP, and the other, Sequenase, which lacks the exonuclease domain.

**Figure 2.**
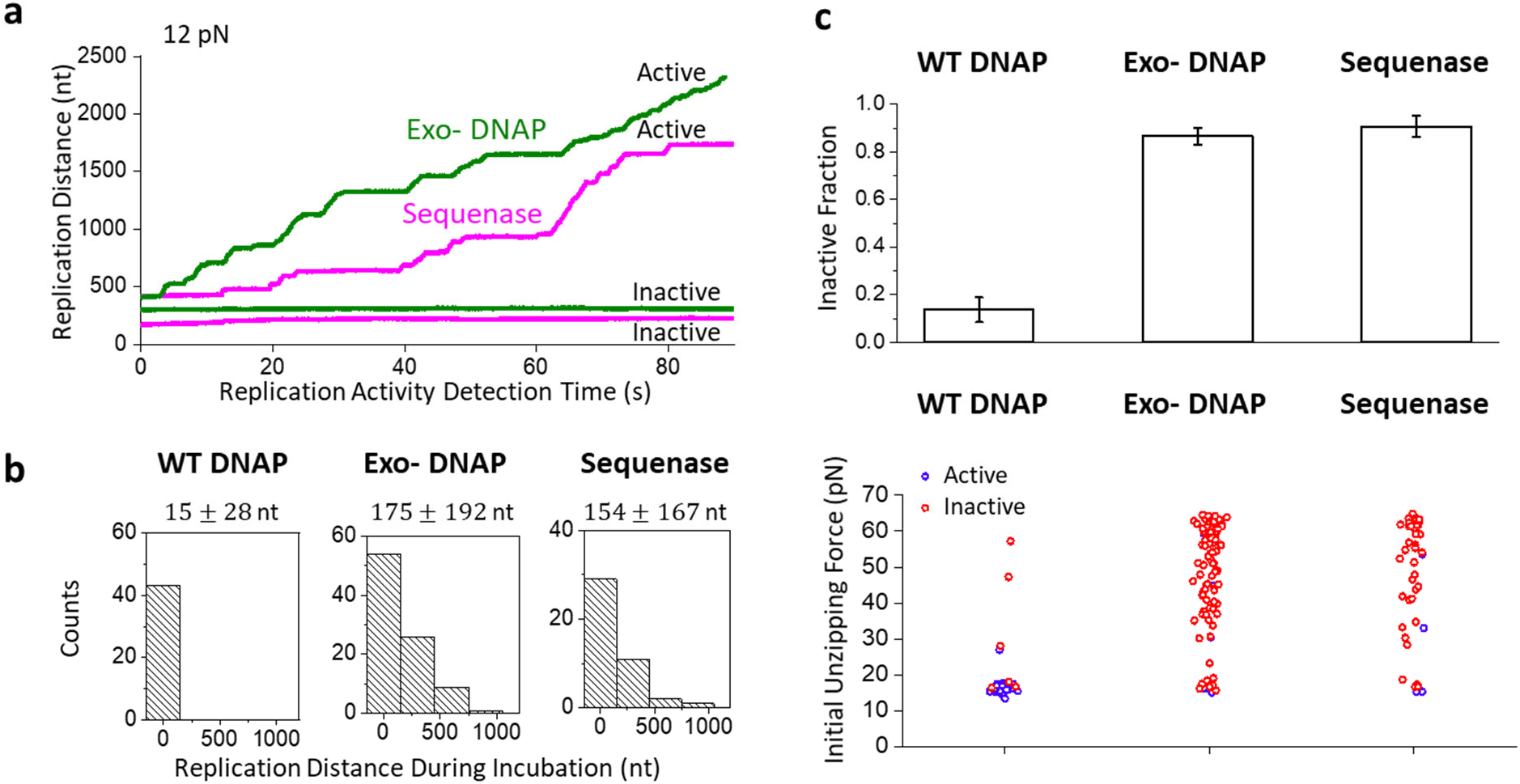
DNAP exonuclease activity limits fork-locking. The unzipping tracker and unzipping mapper techniques are used to investigate the role of the exonuclease activity of DNAP on fork-locking. Two exonuclease-deficient DNAP mutants, exo-DNAP and Sequence, are investigated after a 90 ± 17 min incubation with the fork. **a.**Representative traces of exo-DNAP (green) and Sequenase (magenta) during the unzipping tracker step. **b**. Histograms of the replication distance at the beginning of the unzipping tracker step. The histograms have *N* = 43, 90, and 43 individual traces for WT DNAP, exo-DNAP, and Sequenase, respectively, with the mean and standard deviation indicated. **c**. The inactive fraction and the initial unzipping force of the three DNAPs. Top panel: Error bars represent the standard deviation. The numbers of traces for inactive fractions are the same as in **b**. Bottom panel: The initial unzipping forces have *N* = 34, 87, and 43 individual traces for WT DNAP, exo-DNAP, and Sequenase, respectively. Active forks are colored blue, and inactive forks are colored red.

After incubating the forks with these mutants for 90 min, we detected both active and inactive traces with these two enzymes (Fig. 2a). However, unlike WT DNAP, these enzymes often already replicated many nucleotides before the detection step (Fig. 2b). We found that WT DNAP replicated minimally, about 15 nt (Fig. 2b). In contrast, both exo-DNAP and Sequenase replicated about 170 nt (Fig. 2b). Thus, the inability to move in reverse facilitates the progression of these mutants to move against the fork, even without the assistance of helicase, consistent with previous findings^24-26^.

Significantly, the inactive fraction and the correspondingly locked forks increased dramatically from 15% for the WT DNAP to about 85-90% for the two mutants (Fig. 2c). In fact, the exo-DNAP and Sequenase required much less time to inactivate the fork, reaching this level of inactivity after only 30 minutes incubation (Fig. S4), significantly faster than that of WT DNAP. Our control experiments show that this difference was not due to DNAP degradation during incubation (Fig. S2b). Thus, while the inability to move in reverse facilitated forward movement for these mutants, this inability also made the enzyme more vulnerable to inactivation and fork locking. This finding highlights the important role of exonuclease activity – the ability to move in reverse protects the fork from inactivation.

### DNAP forward translocation promotes fork-locking

The observation that exonuclease activity protects the fork from inactivation suggests that the reverse may also be true: forward translocation could induce fork-locking. To investigate this possibility, we worked with exo-DNAP, which showed significant fork-locking after 90 min incubation. To modulate the polymerization rate, we titrated down the nucleotide concentration. We found that under the 12 pN force, the exo-DNAP replication rate was about 40 nt/s at 1 mM dNTPs and decreased to 10 nt/s at 5 µM dNTPs (Fig. 3a,b).

**Figure 3.**
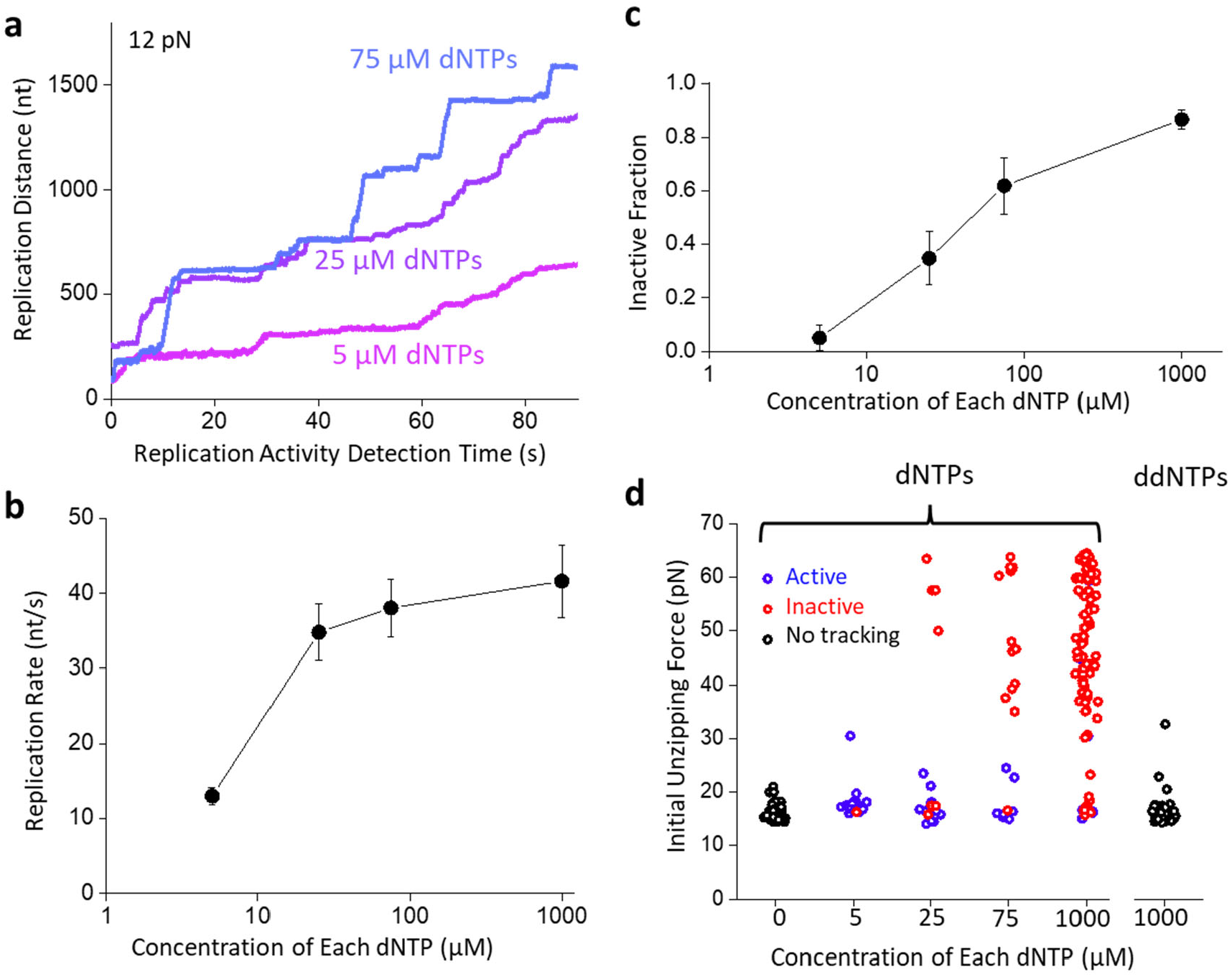
DNAP forward translocation promotes fork-locking. The unzipping tracker and unzipping mapper techniques are used to investigate the role of the forward translocation of DNAP on fork-locking. For these experiments, exo-DNAP was used as it has significant fork-locking activity. **a.**Representative traces of exo-DNAP with reduced dNTP concentrations during the unzipping tracker step. **b**. Replication rate of the exo-DNAP versus dNTP concentration. Data are from *N* = 41, 35, 30, and 20 individual traces with 5, 25, 75, and 1000 µM of each dNTP, respectively. Error bars represent the standard errors of the means. **c**. The inactive fraction of the exo-DNAP at different dNTP concentrations, after a 90 ± 17 min incubation with the fork. Error bars represent the standard deviation. Data are from *N* = 20, 23, 21, and 90 individual traces for 5, 25, 75 and 1000 µM of each dNTP, respectively. **d**. The initial unzipping force of the exo-DNAP at different dNTP concentrations, after a 90 ± 17 min incubation with the fork. Traces that were active during the unzipping tracker are shown in blue, while inactive traces are shown in red, while traces in the absence of dNTPs and with ddNTPS shown in black did not have a tracking step. Data are from *N* = 63, 20, 20, 21, and 87 individual traces for 0, 5, 25, 75, and 1000 µM of each dNTP, respectively. The ddNTP 1000 µM condition contains *N* = 62 individual traces.

Consistent with our prediction, the inactive fraction diminished with a decrease in dNTPs, from 85% at 1 mM dNTPs to 5% at 5 µM dNTPs under 90 min incubation (Fig. 3c,d). There is minimal fork-locking behavior at 5 µM dNTPs. Therefore, these data support the possibility that fork-locking requires DNAP forward translocation. For further confirmation, we replaced the dNTPs with the terminating nucleotides, ddNTPs, which halts replication elongation, and found minimal fork-locking (Fig. 3d).

Collectively, these data show that fork-locking is a consequence of DNAP forward translocation against excessive resistance. Exonuclease activity is essential in minimizing fork locking by allowing DNAP to move in reverse to alleviate the resistance. Without the exonuclease activity, a persistent forward motion of DNAP against a nearly insurmountable obstacle can lead to DNAP locking the fork by interacting with both the leading and lagging strand, rendering the fork inactive.

### Locked replication forks cannot be readily unlocked

The observation that DNAP can lock the replication fork raises the question of whether the fork can be unlocked. We first investigated if DNAP concentration can play a role in this reactivation. We observed a lower inactive fraction and less fork-locking in the presence of 30 nM DNAP versus 1 nM (Fig. 4a). The increase in fork activity suggests that a higher DNAP concentration maintains a more active fork. Inspired by this finding, we investigated whether a higher DNAP concentration could reactivate locked forks. Thus, we incubated the forks under 1 nM DNAP for 90 min, which locked about 15% of the forks, and subsequently introduced 30 nM DNAP to the reaction. We found that the subsequent increase in the DNAP concentration did not reduce the inactive fraction nor the locked fork fraction (Fig. 4a). To further examine this conclusion, we similarly examined exo-DNAP and Sequenase activity (Fig. 4a, Fig. S4).

**Figure 4.**
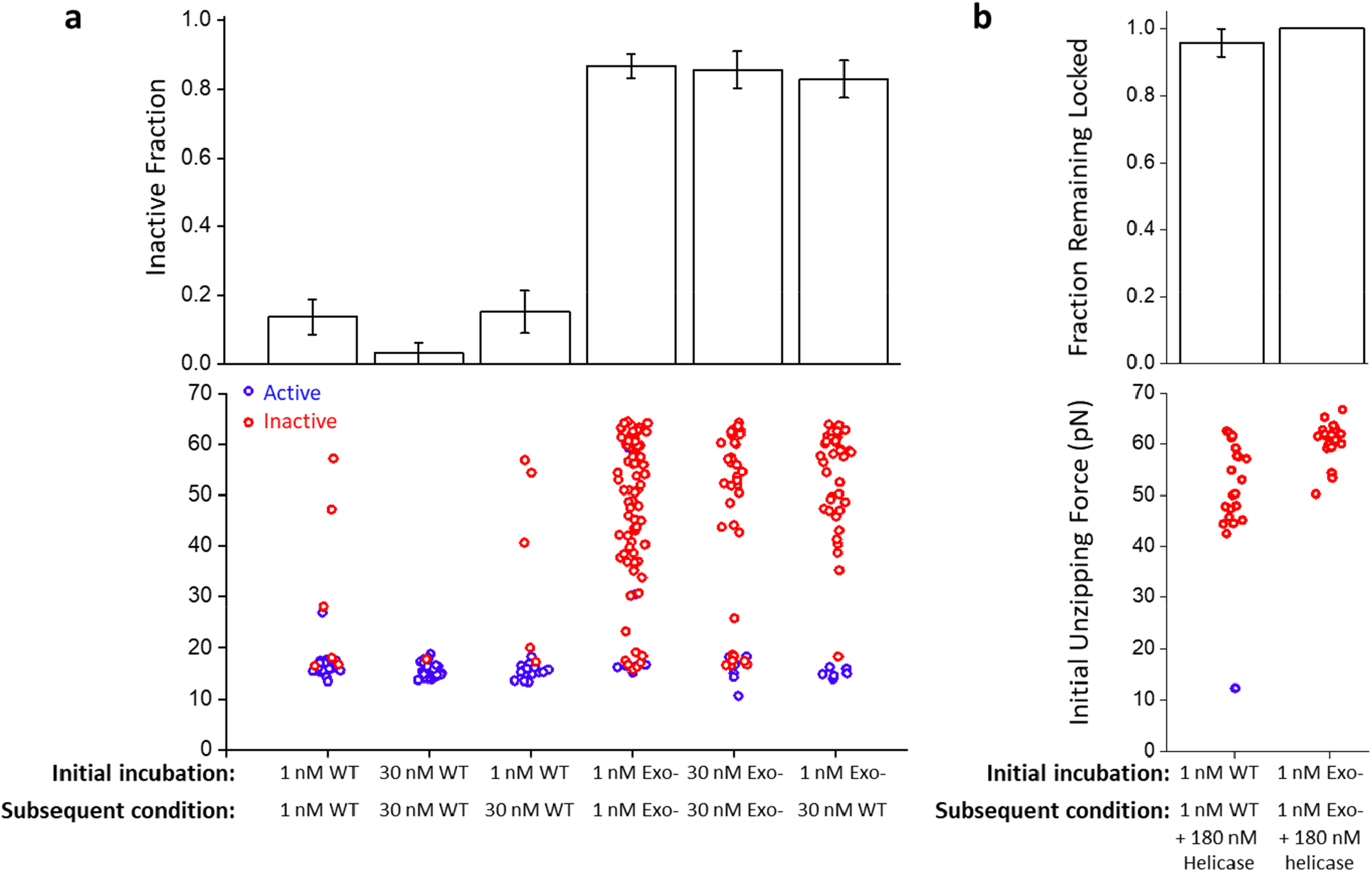
Locked replication forks cannot be readily unlocked. **a**.The inactive fraction and initial unzipping force of WT and exo-DNAP under different protein concentrations. Forks were initially incubated with the indicated DNAP type and concentration for 90 min, and subsequently, the DNAP type and concentration were changed to the indicated subsequent condition. Error bars represent the standard deviations. Inactive fraction traces: *N* = 44, 33, 33, 90, 42, and 48 for conditions shown from left to right. Initial unzipping force traces: *N* = 34, 28, 27, 87, 40, and 47 for conditions shown from left to right **b**. The fraction remaining locked after helicase is introduced to the forks. After the initial incubation of 90 min with WT or exo-DNAP, tethers with locked forks were selected (Methods). After the addition of helicase, these locked forks were checked using unzipping tracker and mapper assays. Reactivated forks are shown in blue, and forks remaining inactive are shown in red. Error bars represent the standard deviations. Fraction remaining locked traces: *N* = 24 and 24 for conditions shown from left to right. Initial unzipping force traces: *N* = 22 and 23 for conditions shown from left to right.

Interestingly, an increased concentration of exo-DNAP and Sequenase made little difference in preventing fork locking, even at a shorter incubation time. When we incubated the forks with 1 nM exo-DNAP, which locked about 85% of the forks, and then subsequently introduced 30 nM WT DNAP to the reaction, we found that the subsequent 30 nM WT DNAP also did not reduce the inactive fraction nor locked fork fraction (Fig. 4a). Taken together, this implies that while excess WT DNAP promotes a more active fork, once the fork becomes locked the excess DNAP cannot facilitate unlocking the fork.

*In vivo*, helicase arrival at the replication fork could rescue a stalled DNAP as helicase’s motor activity may provide the necessary driving force to disrupt the tightly locked fork by DNAP^3, 4^. To investigate this possibility, we slightly revised the experimental approach to improve the data throughput. For this experiment, we used a DNA template with a ssDNA region on the lagging strand to ensure helicase loading (Fig. S1c). We first selected forks locked by WT or exo-DNAP during the initial 90-minute incubation period (Methods). Subsequently, we added helicase to the reaction and examined whether helicase could help unlock these forks. We found that helicase was ineffective at reactivating a locked fork (Fig. 4b). The results indicate that the locking configuration is not readily reversible. This finding highlights the importance of the helicase presence at the fork to keep the DNAP from experiencing excessive resistance. If helicase dissociates from the fork, it must be replaced in a timely manner to maintain an active fork.

## Discussion

In this work, we discovered that T7 DNAP working alone for an extended duration at a replication fork can tightly lock the replication fork by interacting with both the leading and lagging strand. Fork locking is a consequence of forward DNAP translocation and can be minimized via the exonuclease activity of the DNAP which allows DNAP to move in reverse.

Once a fork is locked, it is nearly impossible to reactivate the fork. These findings highlight the importance of the exonuclease activity of the DNAP and the critical need for the presence of helicase at the fork in maintaining an active fork.

*In vivo*, dNTP concentrations are regulated to be low at 5∼37 uM^27^. We found that fork locking is a consequence of DNAP forward translocation and is greatly reduced with reduced dNTP concentrations. We speculate this may provide an evolutionary advantage for replication when dNTP concentrations are low. Under this condition, the helicase may dissociate or slip at the fork^28^, leaving DNAP to work on the fork on its own. The reduced dNTP concentration then minimizes fork locking to maintain an active fork even under sub-optimal growth conditions.

Although our work focuses on the T7 DNAP, our findings may also apply to other prokaryotic DNAPs and eukaryotic DNAPs that replicate the leading strand, such as Pol III of *E. coli* and Pol ε of eukaryotes. It is possible that these DNAPs can also lock the replication fork when working on their own. All of these DNAPs contain an exonuclease domain. Although the most well-established role of their exonuclease domains is proofreading^29-32^, recent studies suggest that the ability to degrade the nascent DNA may promote replication restart^33, 34^. Our findings are consistent with this view and suggest that the exonuclease activity could facilitate fork restart by limiting fork locking and inactivation. In contrast to replicative DNAP, most translesion synthesis (TLS) DNAPs lack the exonuclease function, and usually work in situations in the absence of helicase^35^. It is possible that TLS DNAPs could also lock the replication fork and must be tightly restricted.

This work demonstrates that maintaining a stable and active replication fork is a complex task, even with the relatively simple T7 system used here. The results and experimental approaches presented here may be applicable to all other replication systems and will facilitate future studies into the complex interactions at the replication fork.

## Acknowledgments

We thank members of the Wang Laboratory for helpful discussion and comments. We especially thank Dr. S. Zhang and Biswanath Shaw for some preliminary studies, and T.M. Kay for help with template preparations. This work is supported by the National Institutes of Health grants R01GM136894 (to M.D.W.) and National Institute of General Medical Sciences grant GM118086 (to S.S.P.). M.D.W. is a Howard Hughes Medical Institute investigator.

## Author contribution

X.J., J.T.I, and M.D.W. designed single-molecule assays. X.J. prepared DNA templates. A.S. and S.S.P. purified and characterized T7 gp5 exo- and T7 gp4A’ proteins. X.J. performed single-molecule experiments and analyzed the data. M.D.W. wrote the initial draft. All authors contributed to the manuscript revision. M.D.W. supervised the project.

