## Supplementary Materials for "DNA Polymerase Locks Replication Fork Under Stress"

### Materials and Methods

#### Proteins and Reagents

Wild-type T7 gp5 DNA polymerase (WT DNAP) was purchased from New England Biolabs (NEB, Ipswich, MA). *E. coli* thioredoxin was purchased from Sigma-Aldrich (St. Louis, MO). Exonuclease deficient mutant of T7 gp5 DNA polymerase (exo- DNAP) was purified from *E. coli*[1]. Sequenase Version 2.0 DNA Polymerase (Sequenase) was purchased from ThermoFisher Scientific (Waltham, MA). The T7 helicase gp4A' (T7 helicase) was purified from *E. coli*[2].

Dideoxynucleoside triphosphates (ddNTPs) were purchased from Sigma-Aldrich (St. Louis, MO).

#### DNA substrates

The majority of the optical tweezer (OT) experiments were performed using a DNA unzipping template. [Inman, James T, Nano Letters (2014), Le, Tung T, Cell (2018)]. The template had a "Y" shape, consisting of a ~3.2 kb dsDNA trunk ligated to a pair of ~1.4 kb arms (Fig. S1a). The 3.2 kb dsDNA trunk segment was amplified via PCR from plasmid pRL574 [Karen Adelman, PNAS 2002] with the forward primer ACTGCACCTAGTGATCCGAAGGACAACCTGTTC, and the reverse primer GCTGAGTAACCAGGCTGTAA. The PCR product was digested with the restriction enzyme DraIII-HF (NEB, Cat# R3510S). The arms were amplified from pBR322 Vector (NEB, Cat# N3033S) using a shared forward DNA primer CGCGTTTCGGTGATGACGGTGA, and reverse DNA primers /5BiosG/GTTACGGATCCGCGCTCGGCCCTTCCGG for Arm 1 and /5DiGN/GTTACGGATCCGCGCTCGGCCCTTCCGG for Arm 2. Thus, the end of Arm 1 was labeled with a single biotin tag, and the end of the Arm 2 was labeled with a single digoxygenin tag. The PCR products were digested with the restriction enzyme BsmBI-v2 (NEB, Cat# R0739S) to produce unique DNA overhangs.

To form the fork junction, we used four ssDNA oligos: CAGCGCCAGACTGGGGGCGTCCTGCAGAAGGCTCCCACGACGACACCGAC annealed with /5Phos/GGGAGTCGGTGTCGTCGTGGGAGCCTTCTGCAGGACGCCCCCAGTCTGGCGCTGGCGGTCCCTCTACGCACACGCATCTGGGTCTA to form Adapter 1, and

/5Phos/GGGACAGACGCTGTCCGCGCCAGTGCAGAATAAGGA GTCATTCGTGGGGTGGAC annealed with /5Phos/ACCCAGATGCGTGTGCGTAGAGCGGACCGCGTCCACCCCACGAATGACTCCTTATTCTGCACTG GCGCGGACAGCGTCTG to form Adaptor 2. Adapter 1 was subsequently ligated to Arm 1 and Adapter 2 to Arm 2, followed with a gel purification of each ligation. The purified ligation products were then annealed to form a Y-arm construct. This Y-arm construct was then ligated to the 3.2 kb dsDNA trunk to form the final template. The primers and ssDNA oligos were ordered from Integrated DNA Technologies (Newark, NJ).

For the helicase addition experiments, we used a template almost identical to the template described above, but it contained a 50-nt ssDNA on the lagging strand near the fork (Fig. S1c) to ensure helicase loading. In making this template, the Arm 1 was ligated to the ssDNA oligo (/5Phos/GGGAGTCGGTGTCGTCGTGGGAGCCTTCTGCAGGACGCCCCCAGTCTGGCGCTGGCGGTCCC TCTACGCACACGCATCTGGGTCTA) rather than to Adaptor 1. All other steps remained the same.

**Experimental conditions**

Single molecule tethers were formed in a hydrophobic nitrocellulose-coated grease microfluidic sample chamber [Le, Tung T, Cell (2018)]. The sample chamber was incubated with 20 ng/µl of anti-digoxygenin (Roche, 11333089001) for ~10 minutes, and passivated with 25 mg/mL β-casein (Sigma-Aldrich, Cat# C6905) for ~1 h. Subsequently, ~0.5 pM of a DNA template were incubated in the sample chamber for ~10 min, after which the streptavidin-coated 500-nm polystyrene beads were introduced into the chamber and incubated for ~20 min. All the incubations were performed at room temperature (23°C). The configuration of the formed tethers is depicted in Fig. S1b. After tether formation, the chamber was flushed with replication buffer (50 mM Tris-HCl (pH 7.5), 40 mM NaCl, 1.5 mM EDTA, 10% glycerol, 1 mM dNTPs each, 4mM $MgCl_2$ in excess of the total dNTPs concentration, 2 mM DTT, 0.5 mg/mL β-casein) before flowing in proteins. Unless stated otherwise, all the measurements were conducted in the replication buffer at room temperature (23°C). In the assays where dNTPs concentrations were varied, the $MgCl_2$ was still kept at 4 mM in excess of the total dNTPs concentration.

Unless stated otherwise, the measurements were conducted under the following conditions: (1) wt DNAP 1 nM with 200 nM thioredoxin; (2) exo- DNAP 1 nM with 200 nM thioredoxin; (3) Sequenase DNAP 14 unit/mL with 200 nM thioredoxin; (4) wt DNAP 1 nM and T7 helicase 180 nM (monomer) with 200 nM thioredoxin; (5) Exo- DNAP 1 nM and T7 helicase 180 nM (monomer) with 200 nM thioredoxin. In experiments with varied DNAP concentrations, thioredoxin was kept at 200 nM.

1.8 µM (monomer) of T7 helicase was incubated in replication buffer on ice for ~30 min, to form hexamers, before being mixed with DNAP and diluted into the final concentration.

**Single-molecule experiment methods**

The unzipping tracker and mapper measurements were performed using a DNA unzipping template on a custom-built optical tweezer [Brent D. Brower-Toland, PNAS (2002)]. In the unzipping tracker step, we trapped a polystyrene bead attached to a DNA tether and stretched the arms horizontally to a defined force (Fig. S1b). We then clamped the force for 90 s to monitor replication. Then we applied the unzipping mapper step where we unzipped the remaining template trunk to assess initial unzipping force by moving the coverslip horizontally at 100 nm/s. The force and extension of each tether were recorded during the measurement. Unless otherwise stated, the clamp force was set as 12 pN. We converted the extension to replicated nt as described in the “Data analysis” session below.

For Fig. 4b helicase experiments, tethers were incubated with 1 nM wt or exo- DNAP for 90 min, and stretched up to 30 pN and held for 5 s. Tethers with unlocked forks showed unzipping patterns with a ~15 pN baseline and thus never reached the 30 pN threshold. We marked tethers with locked forks that could sustain a 30 pN force for 5 s by recording their positions. Subsequently, 180 nM helicase with 1 nM corresponding DNAP was introduced into the chamber. These pre-selected tethers with locked forks were measured using the unzipping tracker and mapper methods to check if the forks were reactivated.

**Data analysis**

For the Unzipping Tracker measurement, a trace is considered “active" when the net replication distance during the 12 pN force clamp exceeds 100 bp. For the Unzipping Mapper measurement, the “Initial Unzipping Force” is defined as the max force achieved during unzipping the initial 100 bp.

**Conversion of replication distance**

The DNAP replication distance is determined from the measured force and extension of the DNA tether. As DNAP advances the position of the replication fork by synthesizing dsDNA base-pairs on the leading strand an equal number of nucleotides of ssDNA are released into the daughter strand. Changes in the DNA tether extension are therefore due to an equal number of ssDNA nucleotides and dsDNA base-pairs. The force dependent extension per bp of dsDNA is calculated using the Modified Marko-Siggia worm-like chain [Wang, Michelle, BJ(1997)] and the extension per nt of ssDNA is calculated using the freely-jointed chain model [3].

**Replication velocity algorithm**

The replication velocities shown in Fig. 3b were obtained from the replication traces such as shown in Fig. 3a. The replication activity of each trace is monitored by the unzipping tracker for 90 s. The traces have regions of uniform forward translocation interrupted by pauses. To characterize the pause-free velocity of DNAP forward translocation, we removed the pauses prior to calculating the velocity. To detect pauses, a trace was smoothed with a 0.02s sliding window to generate the dwell time versus position [Karen Adelman, PNAS (2002); Alla Shundrovsky, BJ (2004); Lu Bai, PRL (2007)]. A pause is defined as when dwell time falls below 1.8 s/bp. The pause-free velocity of each trace is a mean of the replication speed of active regions between pauses.

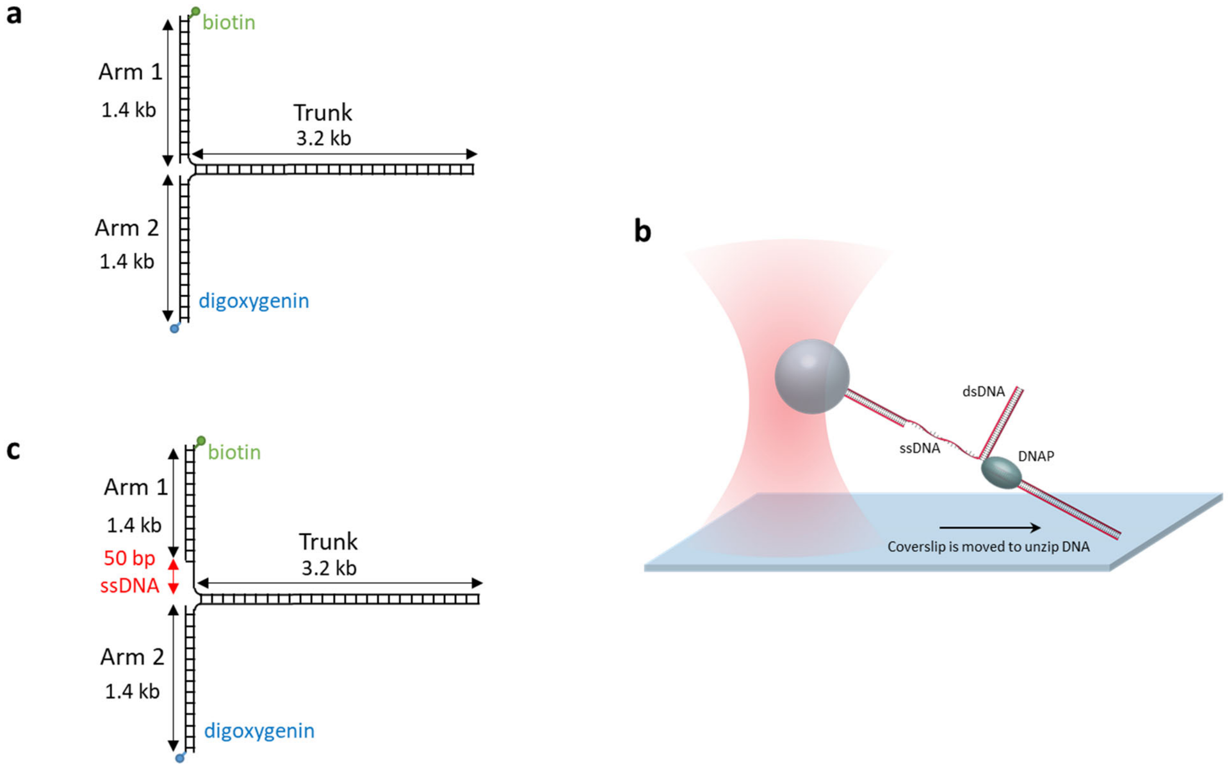


**Fig. S1.** DNA template and setup configuration

**a**. The DNA template used for the unzipping tracker and unzipping mapper experiments.

**b**. Experimental configuration for unzipping tracker and unzipping mapper experiments. The DNA arms formed a tether with one end attached to the coverslip and the other to a polystyrene bead trapped in an optical tweezer. The DNA extension was controlled by moving the coverslip to maintain a constant force (unzipping tracker) or extend at a constant velocity (unzipping mapper).

**c**. The DNA template used for helicase experiment. The ssDNA region was for helicase loading.

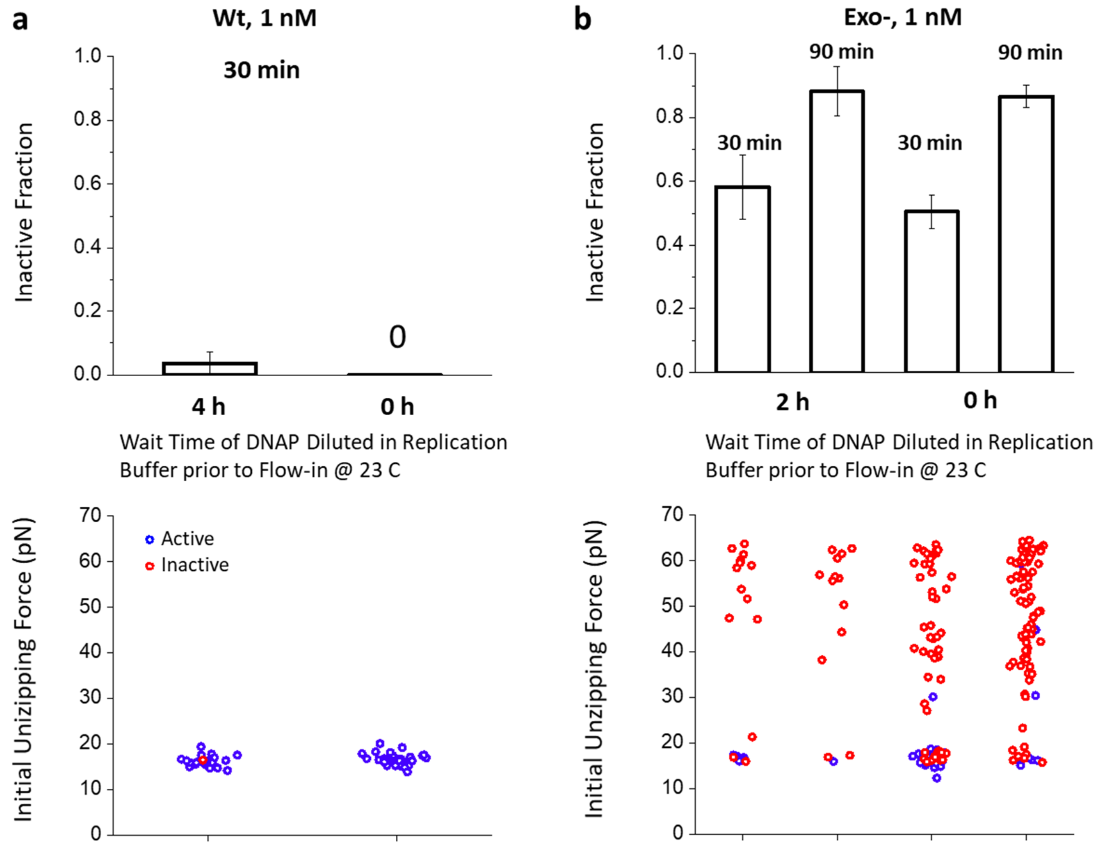


**Fig. S2.** DNAP remains active under single molecule conditions

**a**. The activity of WT DNAP after prolonged incubation under single molecule conditions in the absence of DNA. WT DNAP was diluted to 1 nM in replication buffer and incubated in a test tube at 23 °C for 4 h before introduction into a sample chamber. The activity was tested using unzipping tracker and mapper methods at $30 \pm 17$ min incubation with the DNA fork. The measurement showed minimal inactive or locked forks, similar to the performance of DNAP with no incubation in the test tube. Thus, WT DNAP remained active at 23 °C after 4 h. Error bars represent the standard deviation. Inactive fractions are from N = 27 and 41 individual traces and initial unzipping forces are from N = 22 and 31 individual traces at 4 h and 0 h wait time respectively.

**b**. The activity of exo- DNAP after prolonged incubation under single molecule conditions in the absence of DNA. Exo- DNAP was diluted to 1 nM in replication buffer and incubated in a test tube at 23 °C for 2 h before introduction into a sample chamber. The activity was tested at $30 \pm 17$ and $90 \pm 17$ min incubation with the DNA forks. The fraction of inactive and locked forks remained the same regardless of the time exo- was incubated in the test tube. Thus, exo-DNAP remained active at 23 °C after 2h. Error bars represent the standard deviation. Inactive fractions are from N = 24, 17, 95, 90 individual traces, and initial unzipping forces are from N = 22, 15, 83, 87 individual traces for the conditions presented from left to right.

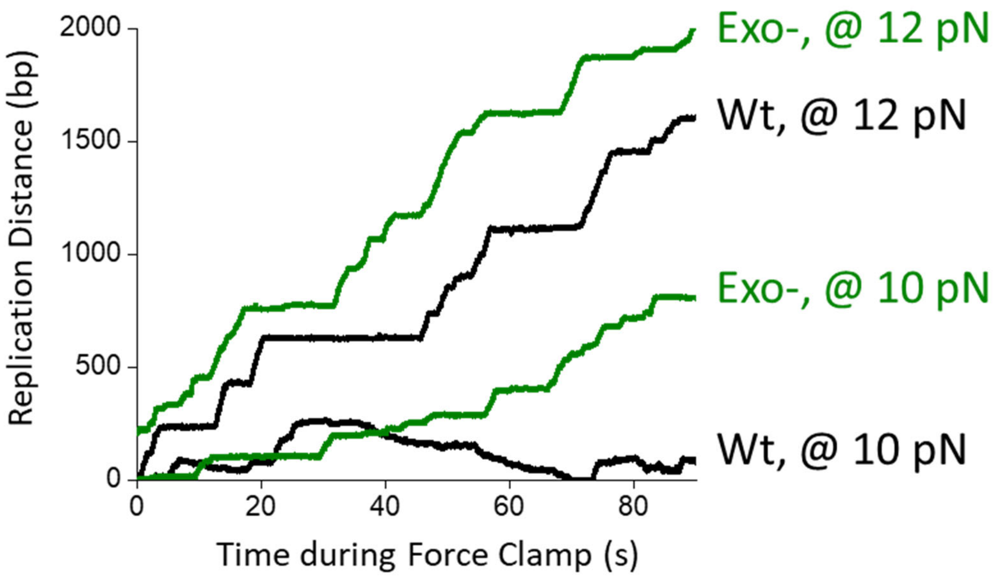


**Fig. S3.** Exo- DNAP had no exonuclease activity

The exo- DNAP was tested for exonuclease deficiency using the unzipping tracker assay. At 12 pN assisting force, Wt DNAP seldom backtracks. However, when we reduced the assisting force to 10 pN, we observed reverse motion of the Wt DNAP which is a result of the exonuclease activity. In contrast, the exo- DNAP only showed forward motion or pauses which confirms the exonuclease deficiency.

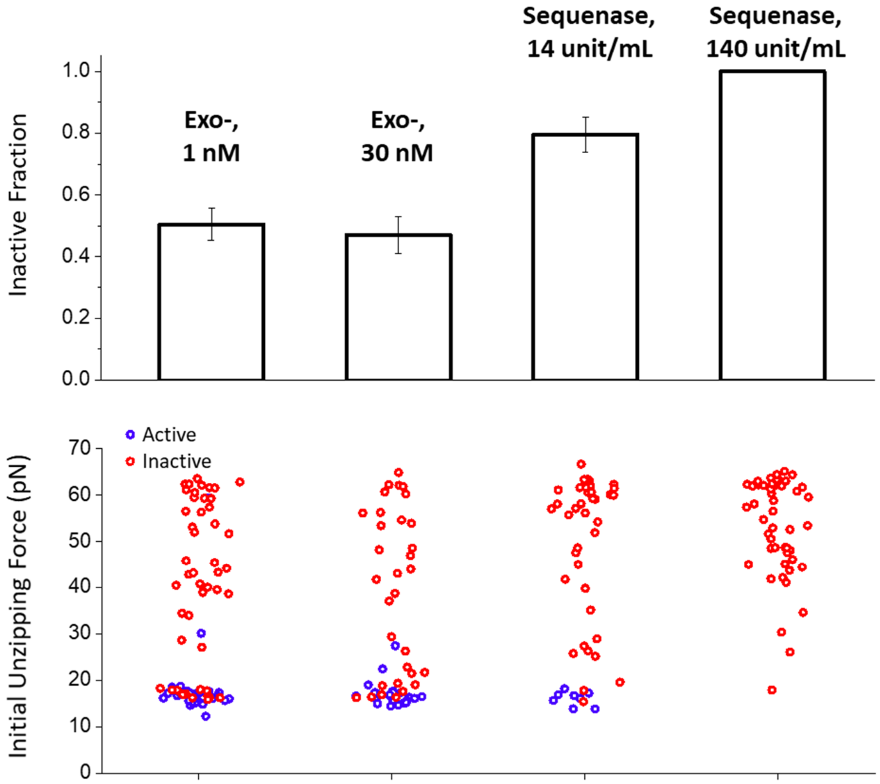


**Fig. S4.** Exo- and Sequenase activity at 30 min incubation and different concentrations

To explore the rate at which exo- DNAP locks the fork, we preformed unzipping tracker and mapper measurements for exo- DNAP and Sequenase at $30 \pm 17$ min incubation. In contrast to Wt DNAP, there is a significant fraction of locked forks after 30 minutes' incubation suggesting that exo- DNAP locks the fork more rapidly than Wt DNAP. For both exo- DNAP and Sequenase, increasing the concentrations made little difference on their inactive fraction and fork locking behavior, strengthening that DNAP exchange of DNAP lacking exonuclease function could not protect forks from being locked. Error bars represent the standard deviation. Inactive fractions are from N = 95, 90, 49, and 49 individual traces, and initial unzipping forces are from N = 83, 87, 49, and 49 individual traces for the conditions presented from left to right.
